# Solitary Silence and Social Sounds: Music influences mental imagery, inducing thoughts of social interactions

**DOI:** 10.1101/2023.06.22.546175

**Authors:** Steffen A. Herff, Gabriele Cecchetti, Petter Ericson, Estefania Cano

## Abstract

The COVID-19 pandemic was accompanied by a marked increase in the use of music listening for self-regulation^1^. During these challenging times, listeners reported they used music ‘to keep them company’^2^; indicating that they may have turned to music for social solace^3^. However, whether this is simply a figure of speech or an empirically observable effect on social thought was previously unclear.

In three experiments, six hundred participants were presented with silence or task-irrelevant music in Italian, Spanish, or Swedish while performing a directed mental-imagery task in which they imagined a journey towards a topographical landmark^4^. To control for a possible effect of vocals on imagined content, the music was presented with or without vocals to the participants, of which half were native speakers and the other half non-speakers of the respective languages.

Music, compared to silence, led to more vivid imagination and changes in imagined content. Specifically, social interaction emerged as a clear thematic cluster in participants’ descriptions of their imagined content through Latent Dirichlet Allocation. Moreover, Bayesian Mixed effects models revealed that music significantly increased imagined social content compared to silence conditions. This effect remained robust irrespective of vocals or language comprehension. Using stable diffusion, we generated visualisations of participants’ imagined content. In a fourth experiment, a new group of participants was able to use these visualisations to differentiate between content imagined during music listening and that of the silence condition, but only when listening to the associated music. Results converge to show that music, indeed, can be good company.

## Introduction

Music can affect thought. It can evoke autobiographical memories^5, 6^, modulate emotions ^7^, and induce mental imagery during unintentional mind-wandering^8^. In particular, mental imagery ^9^ is commonly experienced during music listening. Fifty-seven percent of personal music listening episodes in everyday life is accompanied by mental imagery^10^, 73% of respondents in controlled lab environments report vivid imagery during music listening^11^, and 83% of attendees report that they regularly experience mental imagery during live-concerts^12^. In addition to the prevalence of mental imagery, music can also affect imagined content.

Compared to silence, content imagined during music listening is more vivid and its emotional sentiment can change^4, 13^. The content imagined often forms elaborate narratives and tends to occur at similar time points in the music across listeners^14^. Not only the ‘when’ but also the ‘what’ of music-induced mental imagery shows similarities among listeners^15^. Manual annotations of music-induced imagery reveal that, most commonly, it is visual^16^ and contains scenes of nature or abstract shapes, colours, or objects^17^. Such convergence of imagined narratives during music appears even tighter when listeners share a cultural background^18^.

Furthermore, studies investigating music consumption during COVID-19-related lockdowns reported an increase in time devoted to music listening and reported that music was used for relaxation, mood modulation, escapism, and ‘to keep them company’^2, 3, 19, 20^(for a review see ^21^). These are not isolated instances, as most of the 1868 Spanish citizens sampled in a recent study reported that they valued music ‘a lot’ (58.1%) or ‘often’ (25.7%) for its social company during the lockdowns^2^. These findings suggest that – in addition to affecting memory, modulating emotions, and supporting mental imagery – the thoughts induced during music listening might be related to sociality. This has important implications for evidence-based cognitive therapies that make use of intentional imagination, such as imagery rescripting^22^ and imagery exposure therapy^23^. Should music indeed be shown to systematically induce themes of social interaction into imagined content, then music-based paradigms that induce directed, intentional imagery could be widely used in therapeutic and recreational settings to potentially aid cognitive therapy, alleviate loneliness, and build social confidence.

Thus, we sought to empirically test potential effects of music on imagined social content using a directed mental imagery paradigm (Figure 1) in which participants had to describe quantitative (vividness, imagined time passed, imagined distance travelled) and qualitative experiences (free format description) of their mental imagery after imagining a continuation of a journey towards a landmark. The free format responses where then analysed with natural language processing tools and visualised through stable diffusion. We use an intentional — rather than unintentional — imagery paradigm to facilitate the generalisation of our results to therapeutic settings.

**Figure 1.**
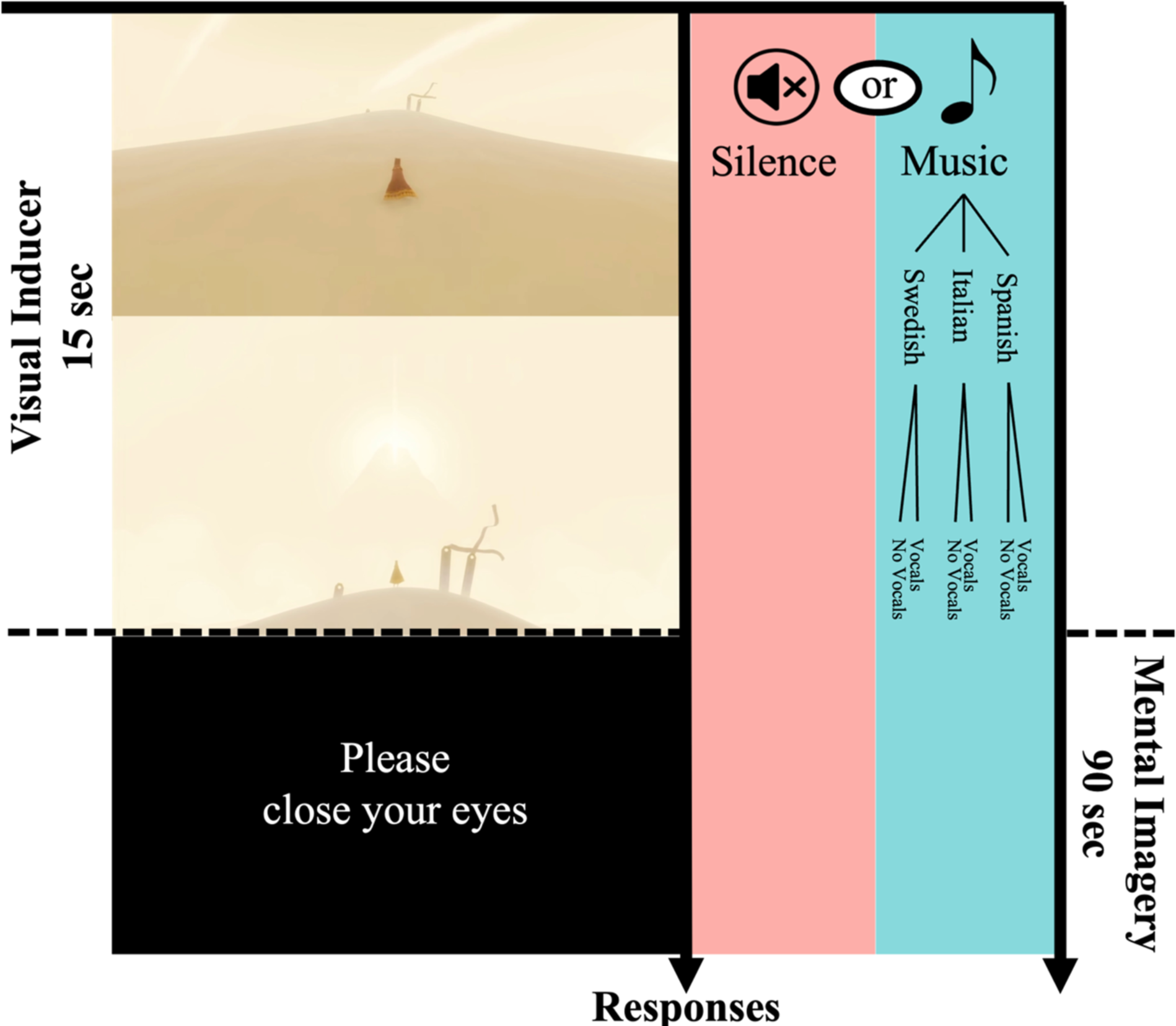
Schematic of the experimental paradigm. Participants viewed a 15-second-long visual inducer video. In the video, a figure ascends a small hill (top left). Once the figure reaches the top of the hill, a large mountain appears in the far distance, barely visible (middle left). Participants then heard a gong and were prompted to close their eyes and imagine a continuation of the figure’s journey towards the mountain. During the 90-second-long mental imagery phase, participants either listened to silence or music. Afterwards, they heard the gong again, prompting them to open their eyes and provide answers to a series of questions about vividness, imagined time passed, imagined distance travelled, and a free format description of their mental imagery before the next trial started. We collected data from 600 participants, split across three experiments using Spanish, Italian, and Swedish songs respectively. In each experiment, we tested both the original songs, as well as versions that had the vocals removed through a source separation algorithm. For each language, we tested 100 native speakers, and 100 non-speakers.

## Results

### Music influences vividness, sentiment, time, and distance imaged

Using Bayesian Mixed effects models, we observed strong evidence (i.e., Evidence Ratios ≥ 39 for undirected hypothesis tests are denotated with ‘*’, see ^24^) that music predicts higher vividness (*β* = 0.46, *EE* = 0.03, *Odds*(*β* > 0) >9999*, Figure 2A), more positive sentiment (*β* = 0.41, *EE* = 0.04, *Odds*(*β* > 0) >9999*, Figure 2B), more imagined time passed (*β* = 0.24, *EE* = 0.04, *Odds*(*β* > 0) >9999*, Figure 2C), and larger imagined distances travelled (*β* = 0.44, *EE* = 0.04, *Odds*(*β* > 0) >9999*, Figure 2D).

**Figure 2.**
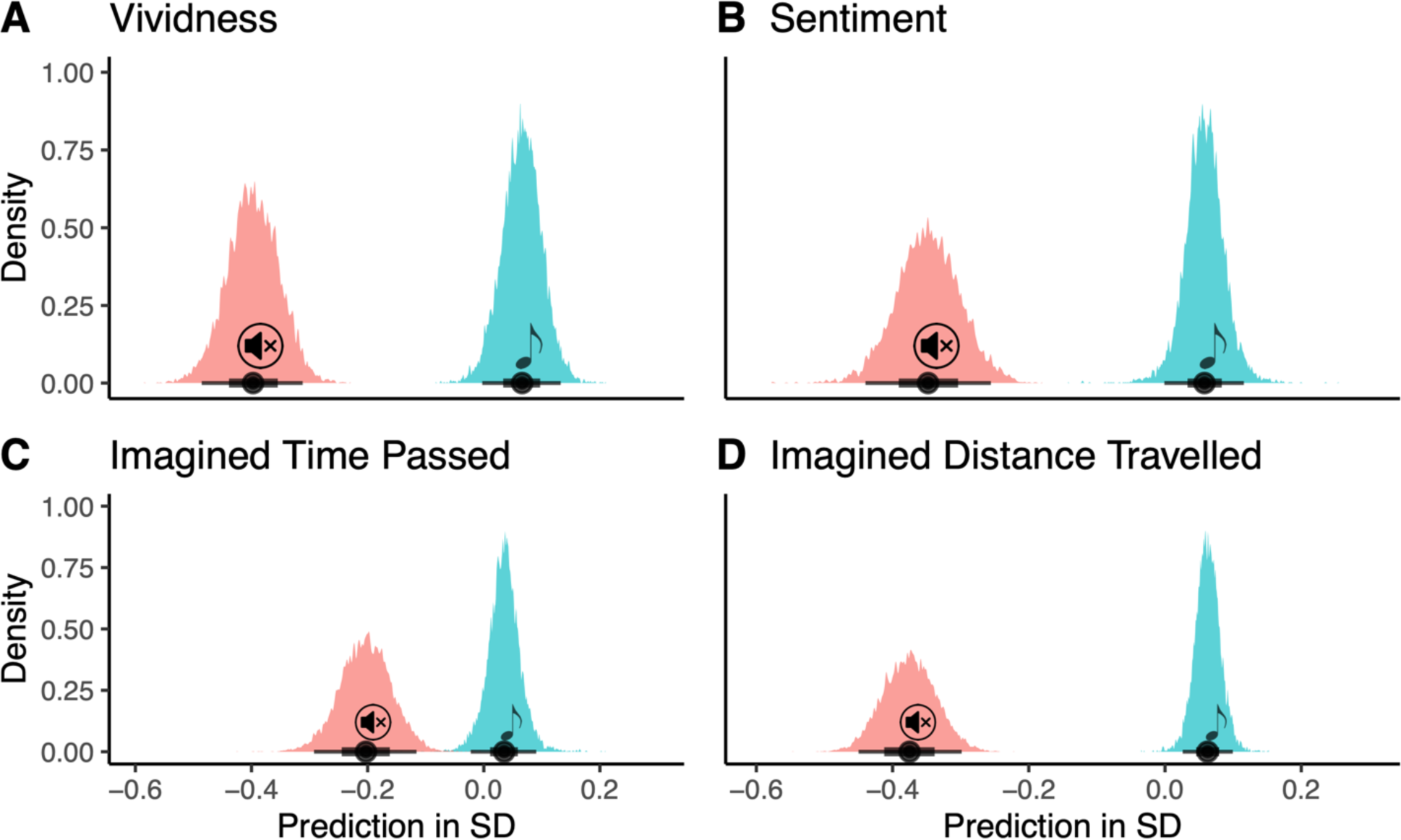
Posteriors distributions of expected predictions for standardised effects for Vividness (A), Sentiment (B), Time passed (C), and Distance travelled (D) in participants’ imagined journeys based on whether participants were listening to music (blue, 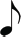) or not (red, 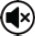).

### Imagined topics differ between silence and music

We analysed the content of the 4200 free format reports of imagined journeys using Latent Dirichlet Allocation^25^ — an unsupervised, bottom-up topic modelling approach to identify latent topic structures— with a nine-topic solution (based on ^26^). For four of the topics (see Figure 3 left, Topics I, II, V, and VI), we obtained strong evidence for differences between music and silent trials.

**Figure 3.**
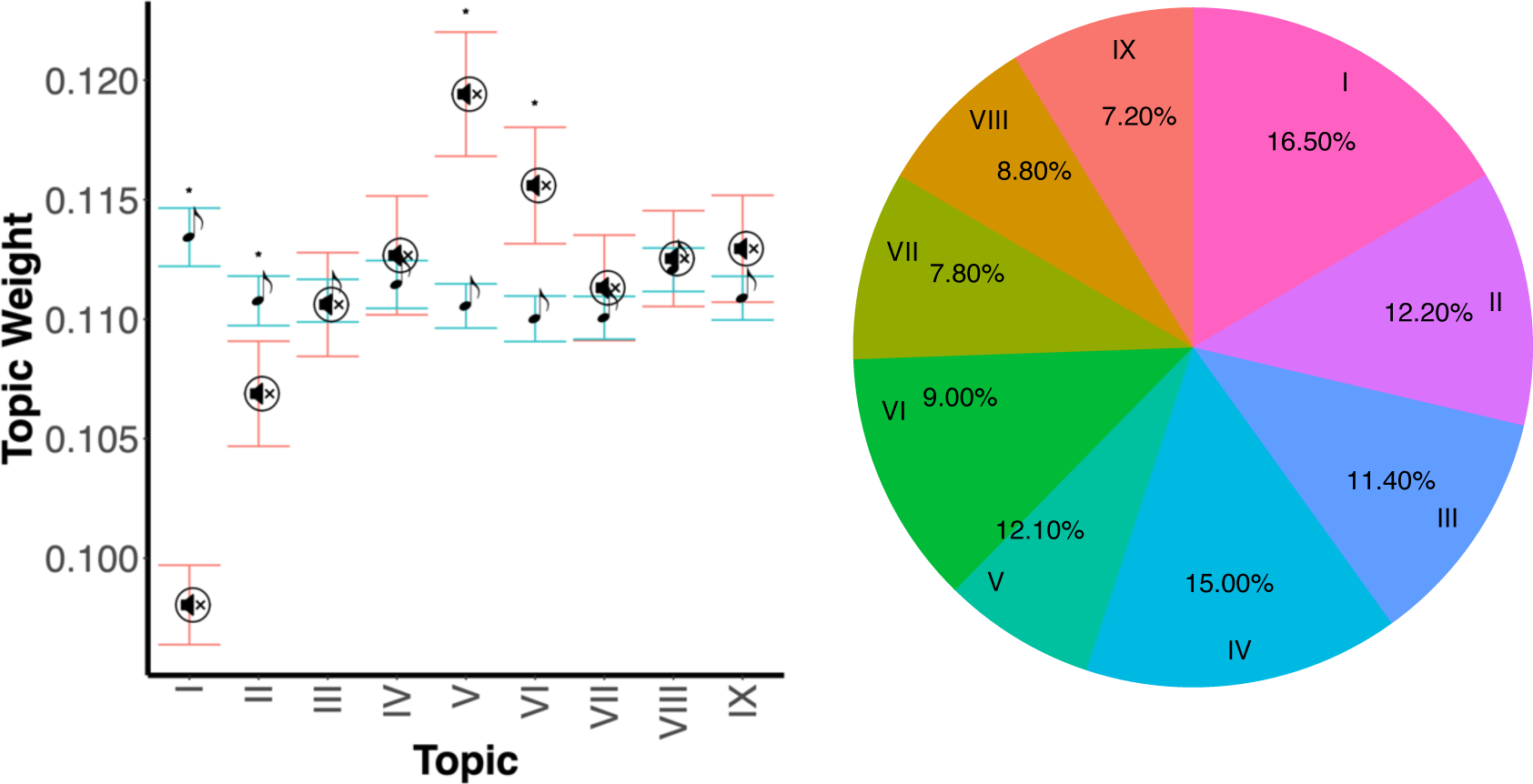
Topic weights by condition (left) and proportion of reports in which the topic was the predominant topic (right). Topics weights I, II, V, and VI differ significantly (*) between participants’ descriptions of their imagined journeys generated during silence (red, 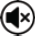) compared to those generated whilst listening to music (blue, 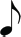). Topic I is the predominant topic most often, and also shows the greatest difference between silent and music trials. Follow-up analysis (see Figure 4) reveals that this topic is associated with themes pertaining to social interactions.

The largest difference was observed in Topic I, a distinct topic containing words related to social interactions (see Figure 4A). We observed strong evidence that music was predictive of an increase in this topic (Topic I, e.g., “*people*”, “*village*”, “*town*”, “*friend*”, *β* = 0.43, *EE* = 0.04, *Odds*(*β* > 0) >9999*). The only other topic which was also more dominant in the music compared to the silence conditions was a topic that contained predominantly words related to heroic adventures, (Topic II, Figure 4B, e.g., “*find*”, “*final*”, “*hope*”, “*fight*”, “sword”, *β* = 0.13, *EE* = 0.04, *Odds*(*β* > 0) = 1999*). On the other hand, two clusters that were more prominent in the silence compared to the music conditions focused on words related to the task context, in terms of general features of the visual inducer video (Topic V, Figure 4C, e.g., “*walk*”, “*sand*”, *β* = −0.31, *EE* = 0.04, *Odds*(*β* < 0) > 9999*) and other aspects explicitly mentioned in the task instructions (Topic VI, Figure 5D, e.g., “*hill*”, “*figure*”, “*gong*”, *β* = −0.19, *EE* = 0.04, *Odds*(*β* < 0) > 9999*).

**Figure 4.**
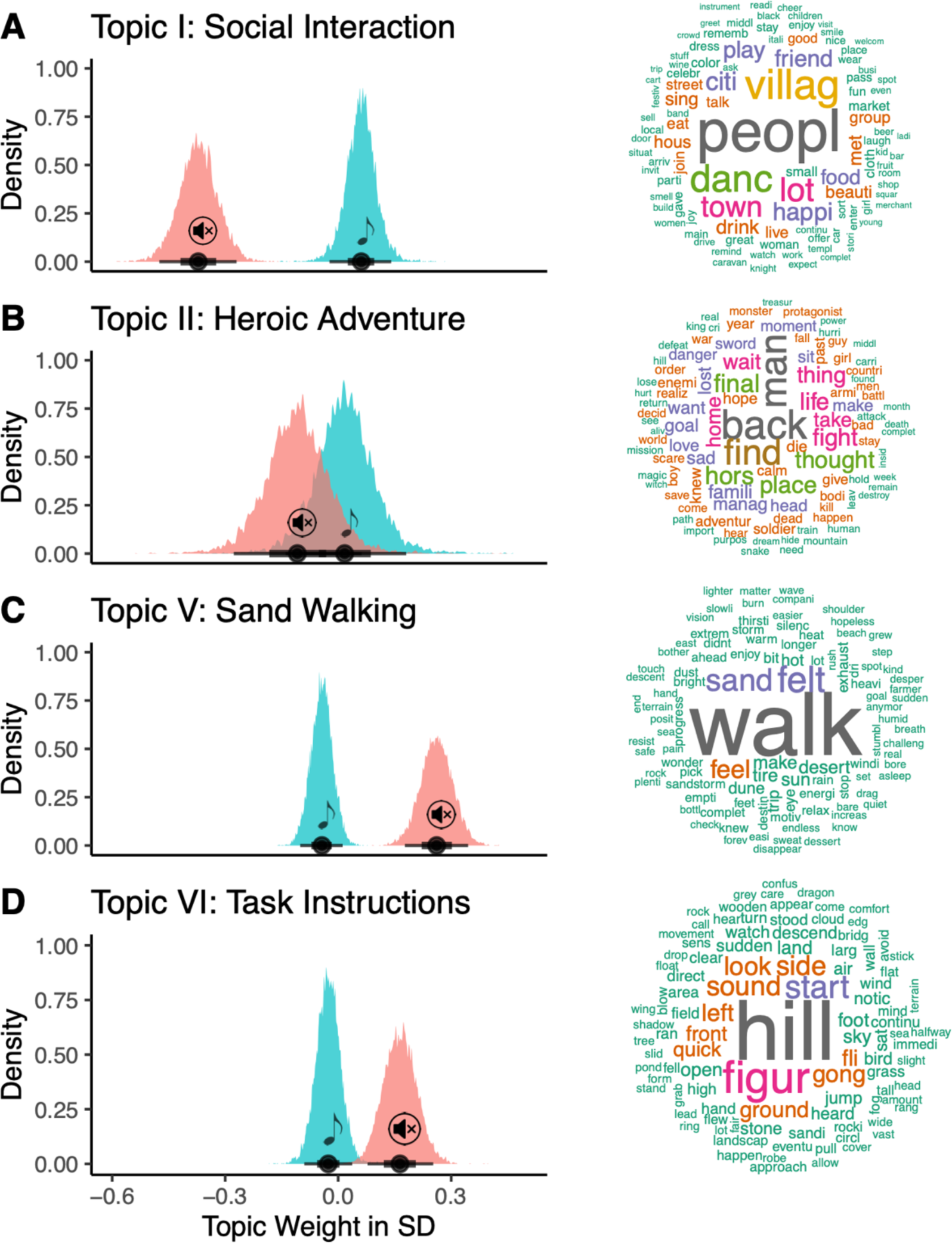
Posterior distribution of the predicted effect of listening condition on the four topics (left) that showed a significant difference between music (red, 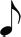) and silence (blue, 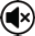), as well as importance weighted word clouds (right), associated with the respective topic. Note, the spelling of some words is off, as all words were lemmatised and stemmed prior to analysis.

**Figure 5.**
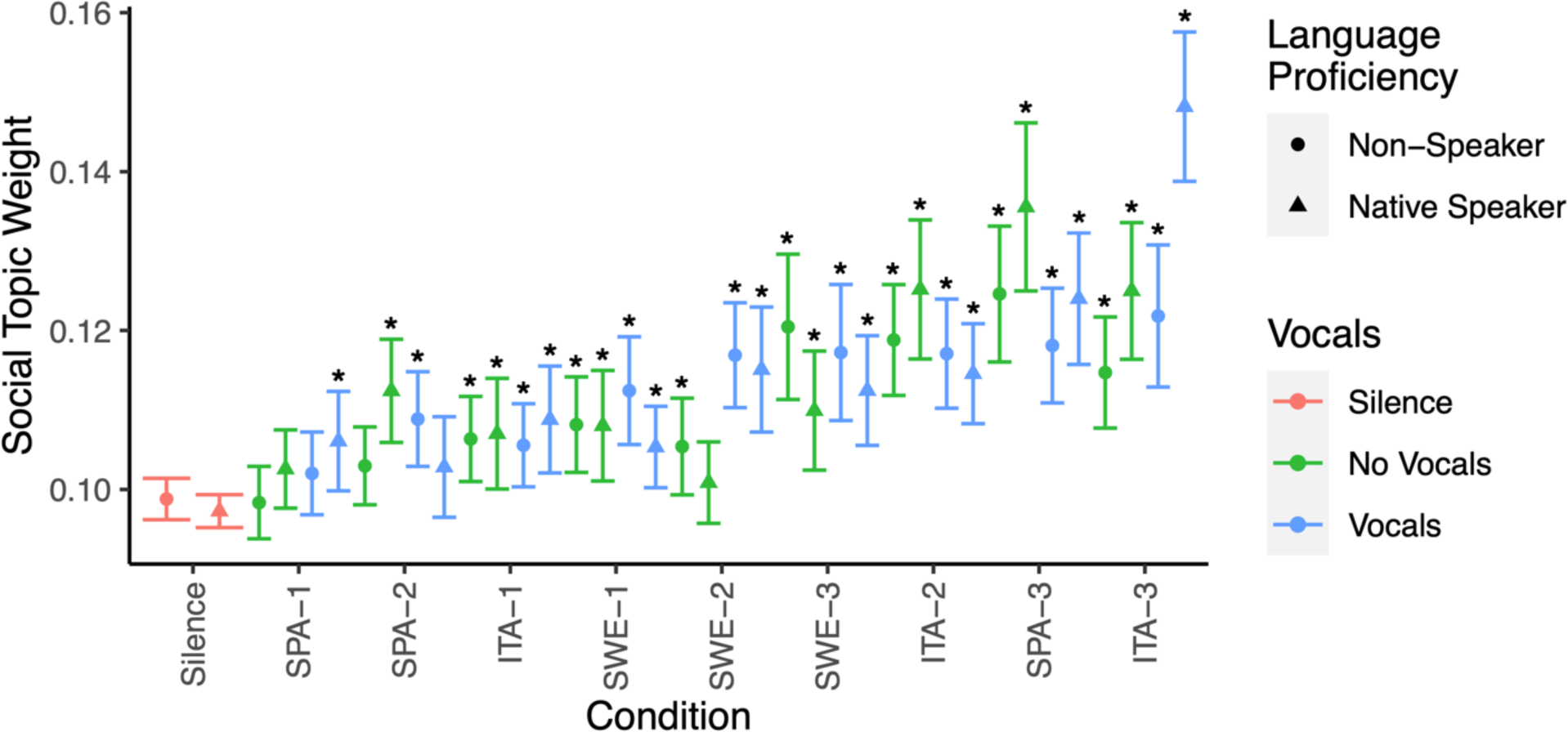
Social topic weight in the imagined content by auditory condition, the presence of vocals, and participants’ language proficiency for the language of the lyrics used in the experiment they are participating in (Spanish, Italian, or Swedish). Compared to silence, the majority (83%) of all music conditions show higher social topic weights in participants’ imagined journeys (indicated by ‘*’).

The increase in the social interaction topic weight was consistent across conditions. In particular, we did not find strong evidence for a general effect of vocals vs. no vocals, nor general effects or interaction based on whether participants were native speakers of the language used in the lyrics of the respective experiment they participated in (all *Odds* < 16.24). Furthermore, strong evidence (i.e., Evidence Ratios ≥ 19 for directed hypothesis tests are denoted with ‘*’ in Figure 5) for higher social topic weights in music compared to silence was observed for 30 out of the 36 music conditions tested (83%). The precise musical piece played seemed to further fine-tune the effect, with some songs showing much higher social topic weights than others (e.g., ITA-3 vs ITA-1, *β* = 1.09, *EE* = 0.13, *Odds*(*β* > 0) = 3999*).

Whilst a detailed musical analysis of all stimuli is beyond the scope of this brief report, ITA-3, in particular, is an interesting case, as it is the only piece showing compelling evidence for an interaction between vocals and language proficiency, whereby the presence of vocals drastically increased the social topic weight, but only if the listener spoke the language (*β* = 0.44, *EE* = 0.18, *Odds*(*β* > 0) = 149.4*). The explanation might well lie in the specific lyrics of the piece (ITA-3: *Nanni ‘Na gita a li castelli’*), which describe social activities around vineyards.

### Musical expertise shows only minimal effects

We did not observe that musical expertise affects or interacts with whether music or silence was present in terms of vividness, sentiment, or the social topic weight of the imagined content (all *Odds* < 11.23). However, we observed that as musical expertise increased, so did the imagined time passed (*β* = .14, *EE* = .04, *Odds*(*β* > 0) > 9999*) and distance travelled (*β* = 0.7, *EE* = .04, *Odds*(*β* > 0) = 33.19*). Intriguingly, this was only observed in the silent but not in the music condition (Time *β* = −0.02, *EE* = .02, *Odds*(*β* > 0) = 0.13; Distance *β* = −.01, *EE* = 0.02, *Odds*(*β* > 0) = 0.2).

### Stable diffusion visualisation of imagined content captures the social topic

To provide readers with an intuitive understanding of the difference between the content imagined in the music and silent conditions, we used a stable diffusion model to visualise participants’ imagined content. Figure 6 (top) shows an example visualisation obtained from a participant’s free-format response in the silent and music conditions. Figure 6 (bottom) shows differences between the silent and music condition based on the latent structure the Latent Dirichlet Allocation identified across all three experiments. For this, the words contained within the nine topics identified by the Latent Dirichlet Allocation and weighted by their respective weights were further adjusted by the specific weight the music or silent condition attached to the respective topic. This means that the bottom row visualises what constitutes a representative, or stereotypical image for the silence and music conditions, based on the Latent Dirichlet Allocation weights. This bottom-up visualisation approach seems to capture the increased prevalence of social interaction across the three experiments.

**Figure 6.**
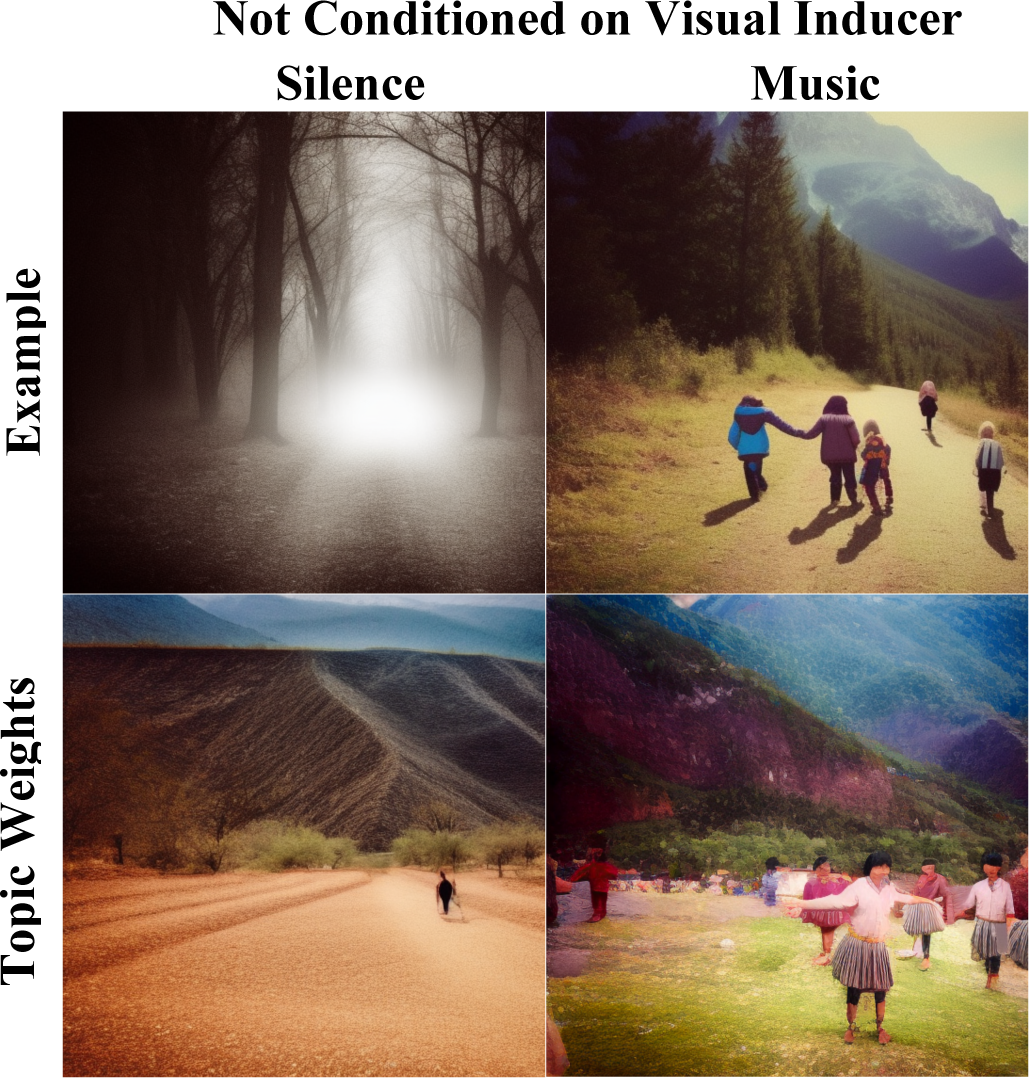
Visualisations obtained through stable diffusion. The top row shows visualisations from one participant’s free-format responses. The participant (ItaS1) described their mental imagery in the silence condition as *‘I imagined a dark walk, without emotions, alone, looking for some hope’*, and in the music condition (ITA-2) as *‘I imagined a walk in the mountains with my family, all together, happy and carefree, we played, we laughed’*. For the images in the bottom row, the stable diffusion model was provided with the average weight distributions across all three experiments, for the respective words of the latent topics, separately for the music and silence conditions. This means that the words and prompt provided to the stable diffusion model in the bottom row are identical between the silence and music visualisation, only the weights for the words differ based on the quantitative results above.

### Not Conditioned on Visual Inducer Silence Music

The stable diffusion primarily serves as intuitive visualisation of our findings and not as inference itself (which we provided above). However, the resulting visualisations do seem to capture both the findings presented above, as well as findings of additional manual annotations of the data set that report generally darker (4% of Music and 12% of Silence trials describe dark settings or the absence of colour), less pleasant temperatures (5% of Music vs. 11% of Silence trials contain description of very hot or very cold temperatures), and less social (39% of Music vs. 12% of Silence trials contain social interactions) imagery in the silence compared to the music conditions. The online supplement (https://osf.io/jgce6/), contains rendered images based on each of the 4200 imaginary journeys, to provide readers with further intuitions of the differences between imagination during music listening and silence, and offer a rich dataset of written and visualised accounts of mental imagery for future studies.

### Participants can differentiate stable diffusion visualisations of content imagined during music and silence, but only when listening to music

To explore whether there is a shared understanding of how music might influence mental imagery of other human listeners, and if so, whether the stable-diffusion images are able to capture this, we conducted one final experiment. Sixty participants who did not participate in the prior three experiments and were sampled from a different population were presented with the stable diffusion images. In each trial, they saw one image generated of the description of the imagined content during the silent condition, and one of a music condition of the same participant. They were then asked to click on the image they thought represented the content the original participant imagined whilst listening to music. Half of the new participants completed the task in silence, whilst the other half completed the task whilst listening to the music associated with the image coming from the music condition. As shown in Figure 7, the task was overall challenging for participants; and we did not observe compelling evidence that participants were able to perform above chance in silence condition (*β* = .07, *EE* = .04, *Odds*(*β* > 0) = 17.95). When participants listened to the original music, however, performance was increased (*β* = .22, *EE* = .06, *Odds*(*β* > 0) = 2665), and we obtained strong evidence that they were able to perform the task (*β* = .29, *EE* = .05, *Odds*(*β* > 0) > 9999*).

**Figure 7.**
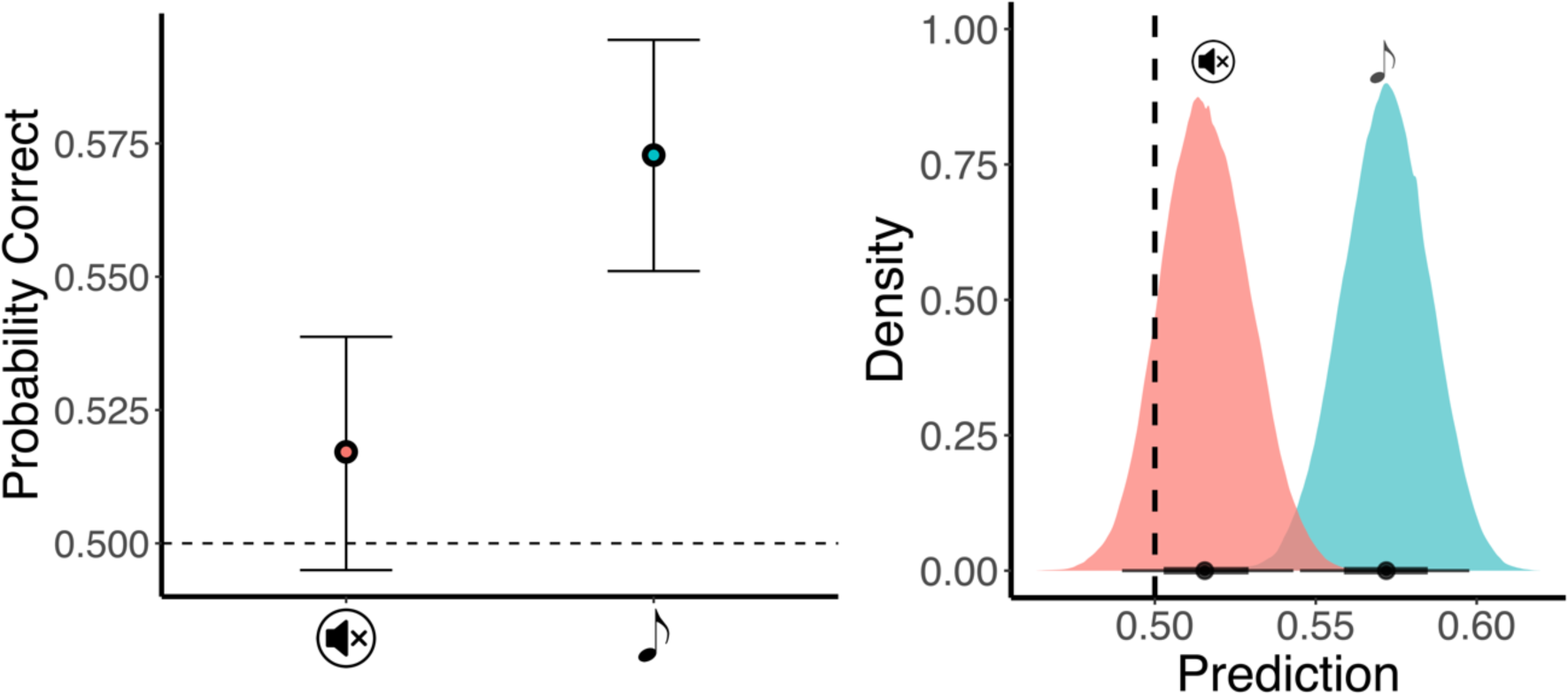
Performance (left) and posterior distribution of expected predictions (right) for differentiating stable diffusion visualisation of content imagined during silence from that imagined during music. The new participants only performed above chance if they were listening to the music (blue, 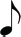) that the original participants heard and not silence (red, 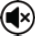) whilst performing the mental imagery task. Error bars indicate 95% CIs.

## Discussion

Many individuals reported turning to music for social solace during the COVID-19 pandemic^2, 3, 19–21^, however, whether this is a figure of speech or an empirically observable effect on social though had not yet been tested. For the first time, we now demonstrate that this is not simply a figure of speech, but that music can indeed induce themes of social interactions into listeners’ thoughts in a directed mental imagery task. Across three experiments on mental imagery, a clear topic related to social interactions emerged bottom-up from the reported imagery of six hundred participants, using an agnostic methodology in the form of Latent Dirichlet Allocation. Prevalence of this social topic was significantly raised in reports of mental imagery obtained when participants listened to music compared to silence. This result was robust and visible in most individual songs tested. Themes of social interactions were increased in music compared to silence, even when the listeners did not speak the language of the lyrics, excluding linguistic engagement and comprehension as an explanation. Furthermore, versions of the songs with the vocals removed entirely did not lead to a systematic decrease in the imagined social interactions, making the presence of the human voice also unlikely as an explanation for the observed effect. This provides empirical credibility and a concrete manifestation to the intuition that music provides social solace, as it does appear to facilitate social thought, which further supports a social function of music^27^.

It is reasonable to assume that music’s ability to induce social interactions into mental imagery is subject to the precise music selected and individual tested. Yet, the observed effects of music seem to generalise relatively well across the experimental conditions. Here, we tested a diverse set of stimuli and participants with varied cultural backgrounds, and we generally found qualitatively darker, colder, and less social imagery in silence compared to music. Furthermore, similar to previous studies, mental imagery during music listening was also more vivid and more positive in sentiment compared to silence^4, 13^. This is of particular interest, as we tested intentional, directed imagery, akin to that utilised in recreational contexts such as roleplay^28^ or in clinical contexts^29, 30^. This highlights music’s potential to support existing and emerging uses of mental imagery. For example, vividness of mental imagery in imagery exposure therapy is a predictor of treatment efficacy^31^ and fine emotional control of imagined content is key in imagery rescripting therapy^22^, and here we show that both –vividness and sentiment– can be affected by task-irrelevant music. It is important to note that music selection here was influenced by the authors familiarity and language proficiency as well as limited by the practical considerations of how many stimuli and participants could realistically be tested. Whilst both songs and participants tested here span a diverse spectrum, generalisability to other musical styles and cultural contexts not tested here –in particular non-Western ones– requires further exploration.

In an additional step, we used stable diffusion to visualise a representation of participants’ mental imagery. This step allowed for the visualisation of aspects of each participant’s individual mental imagery. This also allowed for the generation of a consensus visual representation of content imagined across participants in the two conditions based on the topic weights identified by the Latent Dirichlet Allocation. We propose this as a convenient method to allow researchers and readers to generate an intuitive understanding of the differences between conditions that could easily be applied to existing and upcoming mental imagery studies to complement the statistical analysis. We also showed that participants were able to differentiate above chance between visual representations of mental imagery generated during music listening from those generated during silence. However, this was only the case when participants doing the differentiation task were also listening to the music that imagining participants had previously listened to. This supports previous findings that participants possess some insight of what other participants might imagine whilst listening to music^32^. However, overall, the performance in this task was comparably low. This is likely because this theory of mind of other participants’ music-induced imagined content had to survive two layers of abstraction. First, the abstraction from mental imagery to written responses, followed by the second abstraction in form of the stable diffusion visualisation of the written responses. We consider it remarkable that above chance performance was still possible after these two layers of abstractions paired with inter-individual as well as cultural differences between participants that did the mental imagery task and those that did the matching task. This observation highlights that stable diffusion could be a useful tool to represent participants imagined content.

Future research will have to further investigate the precise underlying musical and acoustic features responsible for music’s ability to affect mental imagery and induce themes of social interactions therein. A thorough understanding of the relevant underlying parameters would enable targeted or even adaptive music composition to affect mental imagery in recreational and clinical settings and would further improve listeners’ access to social solace through music. To this end, we hope that our publicly available data set containing imagery, vividness, sentiment, imagined time travelled, imagined time passed, free format responses, Latent Dirichlet Allocation, topic weights, manual annotations, and corresponding stable diffusion visualisations of all 4200 imagined journeys will prove useful for future research (https://osf.io/jgce6/). Overall, it seems music can indeed be good company.

## Method

### Participants

Data collection of the mental imagery task was split into three experiments, each differing in the stimuli tested (music in Spanish, Italian, or Swedish). In each experiment, we tested 100 native speakers of the respective language, and 100 non-speakers of the language, for a total of 600 participants across the three experiments. Only submissions without missing responses were accepted. All participants were required to be fluent in English and to have normal or corrected-to-normal hearing. We recruited participants through Prolific Academic as it provides access to international samples and offers language proficiency filters. Table 1 provides further descriptive information on each participant sample. Participation was reimbursed with 10 GBP and all participants provided informed consent. The study received ethics approval from the institutional review board of the *École Polytechnique Fédérale de Lausanne* (067-2020/02.10.2020), as well as *Western Sydney University* (H14358).

**Table 1.**
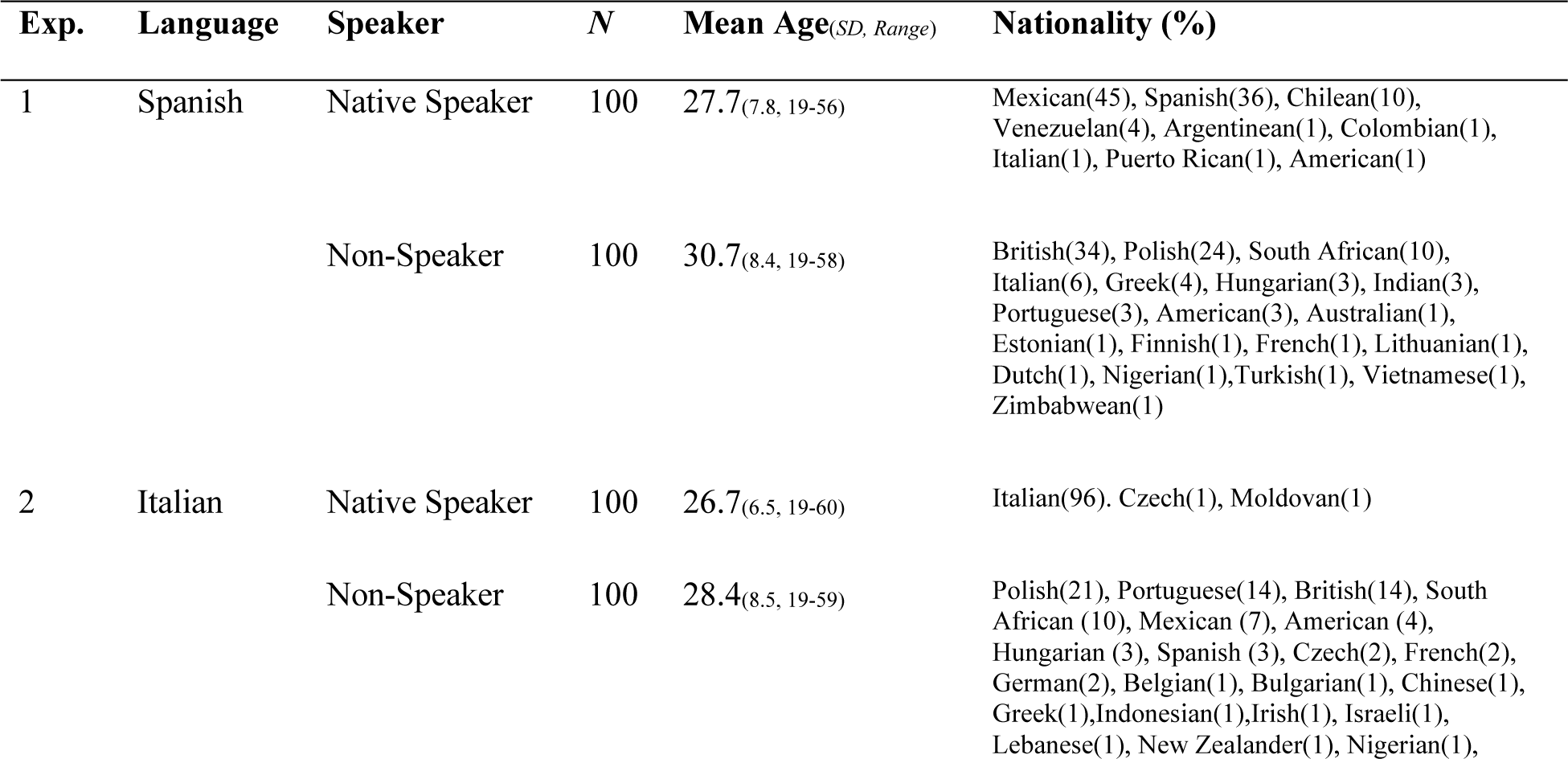
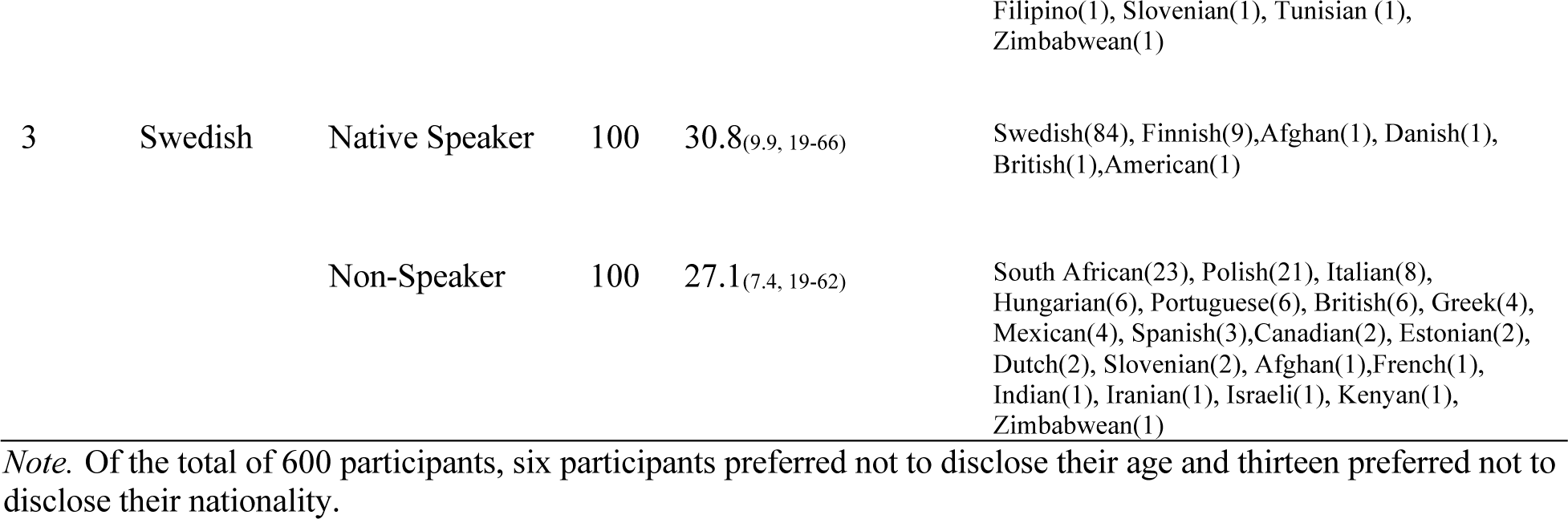
Overview of participant samples.

For the perceptual matching task, we recruited 60 additional participants from the student population at Western Sydney University who were randomly allocated to a music (*N* = 30, *M_age_* = 20.7, *SD_Age_* = 4; self-identified gender: *Female* = 23, *Male* = 6, *Other* = 1) or silence condition (*N* = 30, *M_Age_* = 22.1, *SD_Age_* = 6.6; self-identified gender: *Female* = 27, *Male* = 2, *Other* = 1). None of this sample participated in the original mental imagery task, and Australian participants comprised only a negligible proportion of the participants in the original mental imagery study (0.167%). Participation was reimbursed with course credit. The perceptual matching study also received ethics approval from the institutional review board of *Western Sydney University* (H14358). All participants provided informed consent, and all experiments were conducted in accordance with the declaration of Helsinki.

### Stimuli

All stimuli generally adhered to folk genres with lyrics in Spanish, Italian, or Swedish. We chose these three languages as they are familiar to the researchers conducting this research and participant samples with the respective language proficiency could be recruited in sufficient numbers through the same platform (Prolific Academia). The stimuli for the three studies were selected by the co-author native to the corresponding language. Folk styles across these three experiments vary dramatically, resulting in reasonably diverse set of stimuli (see https://osf.io/jgce6/ for detailed musical feature analyses). The lyrical content of the stimuli also showed great differences, ranging from memento moris to descriptions of grape festivals (see https://osf.io/jgce6/ for original lyrics as well as English translations). The number of stimuli in each experiment was chosen to be compatible with prior research^4, 13^. The first 105 seconds of each song was used to match with the trial duration, starting with the onset of the visual inducer. At the beginning and end of each stimulus, a brief fade-in and fade-out was applied to avoid clipping. Whilst participants could self-adjust loudness to their personal preference, each stimulus was loudness normalized to be comparable with each other and prior studies^4, 13^ to the common value of − 23 ± 5*10^−7^ LUFS, as per EBU R-128 ^33^, using the *pyloudnorm* Python library ^34^.

**Table 2.** Overview of the music stimuli presented to participants in the three experiments alongside a silent control condition.

| Exp. | Language | Song ID | Title | Performer | Composer | Year |
| --- | --- | --- | --- | --- | --- | --- |
| 1 | Spanish | SPA-1 | Preludio para el año 3001 | Amelita Baltar | Astor Piazzola | 1994 |
|  |  | SPA-2 | Luna Tucumana | Mercedes Sosa | Atahualpa Yupanqui | 1993 |
|  |  | SPA-3 | Pedacito de mi vida | Celina Y Reutilio | Celina y Reutilio | 1989 |
| 2 | Italian | ITA-1 | Passacaglia della vita | Marco Beasley | Traditional | 2005 |
|  |  | ITA-2 | Addio mia bella, addio! | Coro e Banda del Tricolore | Carlo Alberto Bosi | 2020 |
|  |  | ITA-3 | Nanni ('Na gita a li castelli') | Lando Fiorini | Franco Silvestri | 1993 |
| 3 | Swedish | SWE-1 | Jämtlandssången | Triakel | Wilhelm Peterson-Berger | 2021 |
|  |  | SWE-2 | Inte har jag pengar inte är jag pank | Ranarim | Traditional | 2006 |
|  |  | SWE-3 | Fager som en ros | Ranarim | Traditional | 2006 |

#### Voice Separation

Each stimulus was presented twice, once with vocals and once without vocals. To create the no-vocals versions, we used a commercial source separation algorithm^35^ on each song. *DemixPro* is a separation algorithm based on deep neural networks trained to separate vocals from backing instruments in music mixes and is generally used for music production applications. The separation process was completely automatic without any pre- or post-processing applied to the tracks or any manual editing. All tracks were in stereo format, sampled at 44.1 kHz.

### Experimental Task

#### Directed Mental Imagery Task

The imagination task deployed here has been previously used to investigate music induced imagination^4, 13^. In this task, participants watched a visual inducer video that serves as common reference for participants’ mental imagery. The video shows a figure travelling up a small hill, before a vague landmark in form of a large mountain appears in the far distance (see Figure 1). The video is a clip from the video game *“Journey”* with written permission from Jenova Chen, CEO of *ThatGameCompany* (see https://thatgamecompany.com).

Fifteen seconds into the video, participants heard a gong sound that prompted them to close their eyes and imagine a continuation of the figure’s journey towards the large mountain. During the 90 seconds of the mental imagery period, the screen is black with white lettering stating, *“Please close your eyes”*. After the mental imagery period, participants heard the gong sound again, prompting them to open their eyes and provide their responses (see Table 3).

**Table 3.** Questions and response format.

| ID | Question | Response Format |
| --- | --- | --- |
| 1 | How much time really passed between the two gong sounds? | Minutes; Seconds |
| 2 | How far away do you estimate the mountain to be at the beginning of the journey? | Kilometres; Metres |
| 3 | How much time passed in your imagination? | Years; Months; Days; Hours; Minutes; Seconds |
| 4 | How far did you travel in your imagination? | Kilometres; Metres |
| 5 | How vivid (clear) was the imagery you experienced compared to experiences in real life? | Numeric response with 0 (not very clear) – 100 (very clear) |
| 6 | Please describe your imagination in as much detail as possible. | Free format text box |

Written instruction on the screen (see https://osf.io/jgce6/ for all instructions) informed participants that if their mental imagery included time and distance skips, then they should include these into their estimates as well. For example, if they imagined a detailed journey of an hour, then imagined travelling for an additional day without filling this journey with distinct content (skipping), and then imagined a continuation for another hour, they were instructed to report 1 day and 2 hours. Questions 1 and 2 only served the purpose of highlighting the difference between real and imagined time as well as distance. As responses differed considerably between these two questions (*Median Time*_|Imagined-Real|_ = 14350sec, *SD* = 21699418, *r* = .012; *Median Distance*_|Imagined-Real|_ = 5000m, *SD* = 228209, *r* = .4) participants seemed to be aware of the difference. Individuals generally differ in the timescales of their imagination. Some imagined journeys on very small timescales (seconds), whereas other participants report differences between journeys on very large timescales (years)^4^. To account for this, question 3 allows participants the opportunity to specify time on a wide variety of metrics.

Each participant was presented with seven trials in random order. Each trial was identical, differing only in whether music was played, and if so, which music it was (see *Stimuli* section). Participants were instructed to *“please treat every repetition independently from one another. You can imagine a similar or a totally different journey every time. This is entirely up to you”* and informed that *“there are no restrictions on your imagination, but please always keep the mountain in sight. This is because after your imagination you will be asked a few questions about the time and distance travelled in your imagination, and the mountain can help you orientate”*.

Upon completion of all mental-imagery trials, participants filled out an online version of the Goldsmiths Musical Sophistication Index^36^. For the present study, we used musical training to account for potential musical expertise effects. In total, the study took 30 to 60 minutes to complete, depending on the depth of detail provided by the participants in question 6.

#### Perceptual Matching Task

Written instructions on the screen explained participants that they would see many pairs of images and that these images were computer-generated based on peoples’ description of what happened during an imaginary journey towards a mountain. They were also informed that during these imaginary journeys, participants were either listening to music or silence. Of the two images shown simultaneously on the screen, one showed content imagined during silence and the other showed content imagined during music listening, based on the descriptions of the same person. After these explanations, participants were instructed that, in each trial, it was their task to click on the picture they thought represented content the person imagined during music listening. After making their choice, a new trial appeared with two images from a different participant. The study was structured in 9 blocks, one for each of the 9 songs of the original mental imagery task. After each block, participants were prompted to take a break and click continue, whenever they felt ready to the start the next block. The order of blocks was randomised between participants, as was the order of trials within each block, as well as the location of the silence and music images within each trial (left or right).

A priori, we established a set of rules to keep this proof-of-concept study conservative. First, we only tested picture dyads where in both trials the original participant was able to generate mental imagery and where the stable diffusion images did not contain any readable text (∼16% removed). This was important as many participants struggled imagining anything during the silent condition and presenting a trial in the matching task, where only one the music-based image showed content, or one of the two images comprises nothing but the written word ‘nothing’ would constitute a confound. Second, we would not correct spelling mistakes before passing the mental imagery description to the stable diffusion model, as this might introduce a non-transparent experimenter bias. Retrospectively, we would advise against this in future studies, as this rule, whilst being more conservative, likely added noise to the data reducing the probability of observing meaningful effects (e.g., some participants reported walking through the dessert rather than the desert…). Third, as the total number of images (4200) was unreasonably large to be presented to each participant and there is a limit to the available student population for the proof-of-concept perceptual matching study, we only tested images generated from the mental imagery of non-speakers in all three experiments, whilst listening to the no-vocals conditions. A priori we reasoned this to be the least homogeneous set of responses and thus likely the most conservative subset available for testing. In each set, participants were randomly allocated to the Music or No-Music group. Participants in the Music condition performed the differentiation task whilst listening to original contextual music that the participants imagining the content also listened to, whereas participants in the No-Music conditions were not provided with the contextual music.

The task took between 30 to 45 minutes, depending on the length of the voluntary breaks between blocks. At the end of the task, participants filled out a demographic questionnaire, provided a subjective difficulty rating of the task (7-point Likert scale, 1 easy to 7 hard), and had the opportunity to share the strategy they deployed to do the task through a free-format text response. All data can be found in the online supplement (https://osf.io/jgce6/).

### Analytical Approach

#### Statistical Inference

Our approach closely followed previous ones^24, 37–44^, including those using the same paradigm^4, 13^. For all statistical inference, we used Bayesian Mixed Effects models^45^, implemented in R^46^ using the brms package^47^. In each model, we standardised all continuous variables (*M* = 0, *SD* = 1), referenced all factorial variables to the silence condition, and provided the model with a weakly informative prior (*t*-distribution; *M* = 0, *SD* = 1, *df* = 3)^48^. In the models for the mental imagery task, the dependent variable (*Vividness, Sentiment, Time, Distance, Topic Weight*, or *manual annotations*) was predicted based on the audio conditions of interest and the participants’ proficiency in the respective language (coded as a Boolean variable), whilst accounting for participant and trial random intercepts. Initially, we also controlled for musical training from the Gold-MSI, however, as it did not carry predictive value in most instances, we dropped this predictor everywhere but where we observed strong evidence in its favour. For the perceptual testing task, we modelled the dependent variable (*Correct Response, Subjective Task Difficulty*) based on the condition (Music vs No-Music), whilst accounting for random effects of participant, trial as well as block number. Instead of a Gaussian identity link as in the previous analysis, we used a Bernoulli logit link to model the binary Correct/Incorrect responses in the perceptual matching task. We report coefficients (*β*), the estimate error of the coefficient (*EE)*, as well as the evidence ratios for a hypothesis test that a given predictor has an effect greater or smaller than 0 (*Odds(β > or < 0)*). Like previous studies, for convenience we use ‘*’ to denote effects that can be considered ‘significant’ under an alpha level 5% (i.e., evidence ratio ≥ 19 for one-sided, and ≥ 39 for two-sided hypothesis tests, see ^24^).

Vividness ratings were provided by the participants responses. Sentiment was calculated by first using the *python* library NLTK^49^ to filter stop-words, lemmatize, and stem participants’ free format response. Then, we estimated sentiment on the pre-processed data with the VADER library^50^, which uses its reference dictionary to map lexical features to an intensity continuum from emotionally negative to positive. We then averaged the word-wise sentiment scores for each report of each participant, to obtain one score per trial. Whilst directly provided by the participant, imagined time and distance varied dramatically between individuals, ranging from

0.1 meters to over 1 million kilometres and 1 second to more than 27 years. As this study focuses on the variations of imagined time and distance across conditions, we log-scaled reported times and distances, and then standardised them (*M* = 0, *SD* = 1) for each participant.

#### Topic Analysis

We first pre-processed the free-format responses by removing stop-words, punctuation, numbers, words that occur fewer than 5 times across the 4200 reports, and extra white space, and then converted all letters to lower case and stemmed all words. We then deployed Latent Dirichlet Allocation^25^ using the *LDA* package^51^ in R, an unsupervised method to identify the content of a fixed number of topics within the corpus. To identify the number of topics prior to analysis, as seen in Figure 8, we used a common density-based metric^26^ implemented in the *ldatuning* package^52^.

**Figure 8.**
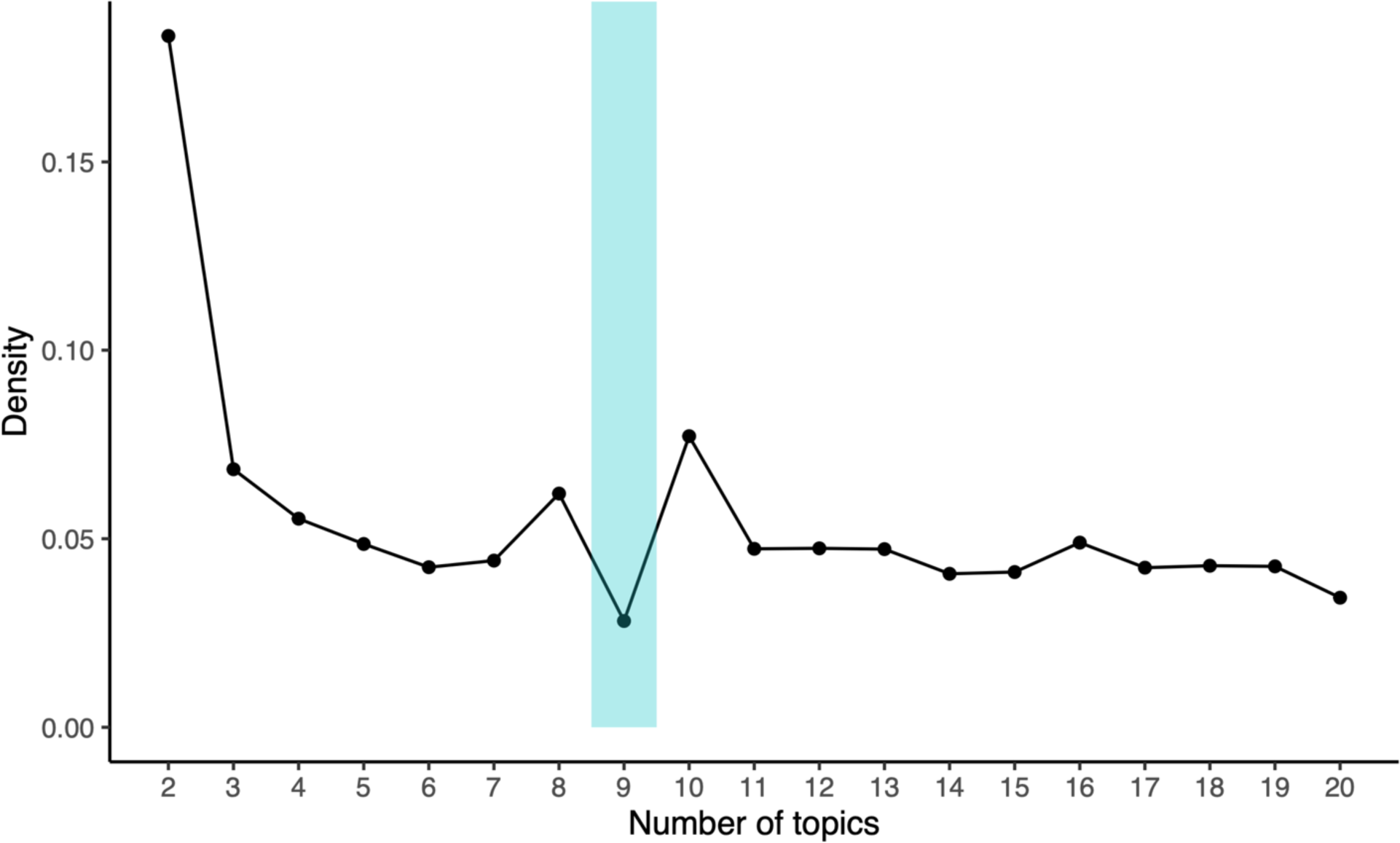
Number of topic search. The topic model performs best when the density (based on average cosine distance between topics, see ^26^) reaches a minimum. Based on this analysis, we set fixed the number of topics to 9 (blue shade) prior to investigating the individual topics.

#### Stable Diffusion Visualisation

We used the *Automatic1111 stable diffusion web UI* (https://github.com/AUTOMATIC1111/stable-diffusion-webui), based on the *Gradio* library^53^, accessed through custom Python code (based on https://github.com/TheLastBen/fast-stable-diffusion, also see *Dreambooth*^54^. The main reason why we opted for the *Automatic1111 stable diffusion web UI* is because it allows to weigh each input word separately. Whilst for the visualisation of individual reports (see https://osf.io/jgce6/), as exemplified in Figure 6 top row, equal word weighting is sufficient, making most stable diffusion models a viable choice, this becomes more complex for the aggregated images as seen in Figure 6 bottom row. For these, we made use of the Latent Dirichlet Allocations’ 9-topic solution. Each topic consists of numerous words associated with this topic, as well as the relative weight of each word to this specific topic (for example, see Figure 4). For the images seen in the bottom row of Figure 6, we first identified in each topic the five words with the strongest weights, and we standardised the corresponding weights by dividing the weights by the maximum weight observed within the respective topic to avoid punishing topics with flatter word weights (e.g., topic II) or prioritising topics with more spiky distributions (e.g., topic I or V) during visualisation. Then, we min-max scaled the relative average importance of the different topics (.09 to .12 on the LDA’s scale, see Figure 3), separately for the music and silence condition, to achieve a more effective weighting scale for the stable diffusion model (0.2 to 2). Finally, we multiplied the standardised weights of the words contained in each topic with the respective scaled topic weight in the music and silence condition separately. This results in two word lists with word weightings that we can use as visualisation prompts for the stable diffusion mode, whereby the silence and music visualisation contain exactly the same words (as determined by the 9-topic solution) but differ in the weights attached to them, based on the data observed in the three experiments.

In a separate step, in order to generate images that resembled the visual inducer more closely, we provided the model with a seed image from the video that participants saw (first frame, shown in Figure 1, top left). As stable diffusion can differ based on the random seed, all the examples shown in Figure 6 were generated with a fixed random seed 14358067202002102272, i.e., our concatenated ethics approval number). Note that we are not claiming that these images show what participants visualised to any particular degree of approximation; instead, we provide these images as an intuitive visualisation of the difference between the music and silence condition both on an individual level (e.g., Figure 6 top row) as well as while abstracting summary features at the population level (Figure 6 bottom row) level. These visualisations may also serve as objects for further research.

#### Manual Annotations

To further corroborate our bottom-up, data driven analysis, a research assistant —unfamiliar with the study’s design and aims — annotated all mental imagery reports in terms of imagined social interactions, temperature, brightness, and narrative perspective (69% First: *‘I’*; 27% Third: *‘He’, ‘She’, ‘It’, ‘They’;* 4% Unclear, similar between Music and Silence trials). The annotator was only provided with the free format accounts of the imagined journeys and did not have access to any other aspect of the data, such as experimental conditions, other responses, or demographics. The exact instructions and definitions given to the annotator can also be found in the online supplement (https://osf.io/jgce6/).

## Acknowledgements

This work was supported by the Swiss National Science Foundation (SNF) under the SPARK grant scheme awarded to Dr. Steffen A. Herff (CRSK-1_196567 / 1) and by the Australian Government through the Australian Research Council (ARC) under the Discovery Early Career Researcher Award (DECRA) awarded to Dr. Steffen A. Herff (DE220100961). We thank the members of the MARCS Music & Science group, in particular Prof. Roger Dean, Dr. Andrew Milne, and Uğur Kaya, as well as Prof. Christian Herff, Dr. Felix Dobrowohl, and Dr. Lauren Fairley for constructive feedback on an earlier draft of the manuscript. We thank Farrah Sa’adullah for her support doing the data collection of the perceptual matching task. Finally, we thank all our participants who shared their imagination with us.

